# Alveolar metabolite availability facilitates secondary infection by *Pseudomonas aeruginosa* in acutely injured lungs

**DOI:** 10.1101/2024.12.18.629020

**Authors:** Jennifer M. Baker, Thomas L. Flott, Laura A. McLellan, Ingrid G. Bustos, Jose L. Guerrero, Lina M. Mendez, Annastasia M. Petouhoff, Piyush Ranjan, Joseph D. Metcalf, Roderick A. McDonald, Nicole R. Falkowski, Ying He, Mónica P. Cala, Gary B. Huffnagle, Michael W. Sjoding, Luis F. Reyes, Kathleen A. Stringer, Robert P. Dickson

## Abstract

Secondary bacterial pneumonia is a frequent complication of acute lung injury (ALI) and the acute respiratory distress syndrome (ARDS). Prior efforts to explain increased pneumonia risk in ALI/ARDS have focused on impaired immunity and bacterial virulence, but have overlooked the potential contribution of ecological factors within acutely injured lungs. Here, we show that lung injury profoundly alters the alveolar metabolic microenvironment, a change that can be exploited by a common pneumonia pathogen. In mice and humans, we found that growth of *Pseudomonas aeruginosa* (the most prominent respiratory pathogen in a retrospective cohort of patients with ARDS) is enhanced by increased alveolar concentrations of multiple ALI/ARDS-associated metabolites, which are derived from the blood and cross the compromised alveolar-capillary barrier during alveolar leak. Collectively, this work reveals an ecological mechanism by which pathogens may survive in the injured lung microenvironment and identifies new potential targets for secondary pneumonia prevention and treatment.

## Main

Secondary bacterial pneumonia is a frequent complication of acute lung injury (ALI) and the acute respiratory distress syndrome (ARDS) and contributes to poor clinical outcomes via increased mechanical ventilation duration,^1–3^ hospital stay duration,^4^ and healthcare costs.^5^ Across clinical observations and animal models, acutely injured lungs are more susceptible to bacterial infection. In humans, secondary bacterial infection occurs across ALI/ARDS etiologies, including severe viral infection^6–8^ and non-viral causes of ARDS.^9^ These clinical findings have been recapitulated experimentally using a variety of animal models that combine bacterial pneumonia with ALI/ARDS-inducing insults such as barotrauma,^10,11^ hyperoxia,^12,13^ or influenza infection.^14^

Prior efforts to understand the increased susceptibility of injured lungs to infection have focused on bacterial virulence^15^ and impaired host immunity.^16^ Yet, research on secondary pneumonia pathogenesis has not accounted for recent insights into respiratory tract microbial ecology.^17^ Once considered sterile, the mammalian lung is now known to harbor dynamic, low-biomass bacterial communities.^18^ In health, the lung microbiome is shaped by bacterial immigration via microaspiration and elimination via local immune defenses and limited nutrient availability.^19^ During ALI/ARDS, however, lung bacterial communities are profoundly altered,^20^ including enrichment with pneumonia-associated^1,21^ and gut-derived bacteria.^22^ Such community-level changes in lung microbiota are detectable with culture-independent methods, predict patient outcomes, and occur even in patients without clinical evidence of pneumonia.^23,24^ These results thus suggest that *ecological factors* (*e.g.*, nutrient availability, oxygen tension, and temperature) likely play a role in the pathogenesis of secondary pneumonia within the alveolar microenvironment.

Thus, to interrogate ecological factors that contribute to secondary bacterial pneumonia in acutely injured lungs, we conducted a series of translational studies to determine the relationship between the growth of *P. aeruginosa* (a frequent etiological agent of secondary pneumonia) and nutrient availability within the acutely injured lung microenvironment. In a retrospective cohort, we found that, consistent with prior reports, rates of ventilator-associated pneumonia (VAP) caused by *P. aeruginosa* were increased in patients with ALI/ARDS. In a murine model of ALI/ARDS, we found that blood-derived metabolites leak into the alveolar space via pulmonary edema, promoting *P. aeruginosa* growth via increased nutrient availability. We validated our murine findings in a prospective human cohort by demonstrating that *P. aeruginosa* growth is enhanced in bronchoalveolar lavage fluid (BALF) from patients with ALI/ARDS and that human BALF metabolite enrichment is positively correlated with both alveolar leak and *P. aeruginosa* growth. These data provide evidence that enhanced alveolar metabolite availability via pulmonary edema is an important ecological factor that contributes to secondary bacterial pneumonia in acutely injured lungs.

## Results

### P. aeruginosa is a major cause of secondary bacterial pneumonia in patients with ARDS

We first investigated the frequency and etiology of secondary bacterial pneumonia in mechanically ventilated patients with ARDS compared to those without ARDS. To do this, we interrogated the electronic medical record of the University of Michigan hospital system and identified 783 patients who received invasive mechanical ventilation for ≥ 48 hours (**Fig. 1a**). Of these, 258 patients met criteria for ARDS^25^ and 525 patients did not (**Extended Data Table 1**). Within this cohort, the daily incidence of VAP in the ARDS group was nearly twice that of the non-ARDS group (**Fig. 1b**). Consistent with prior studies,^26^ *Staphylococcus aureus* and *P. aeruginosa* were the two most commonly identified pathogens (**Extended Data Table 2**). The relative frequency of positive cultures yielding *P. aeruginosa* was greater (*p* = 0.04) than that of *S. aureus* among ARDS patients but not among non-ARDS patients (**Fig. 1c**). We thus concluded that consistent with previous reports, secondary bacterial pneumonia occurs with increased frequency in patients with ARDS, and ARDS patients have increased susceptibility to *P. aeruginosa* infection.

**Figure 1.**
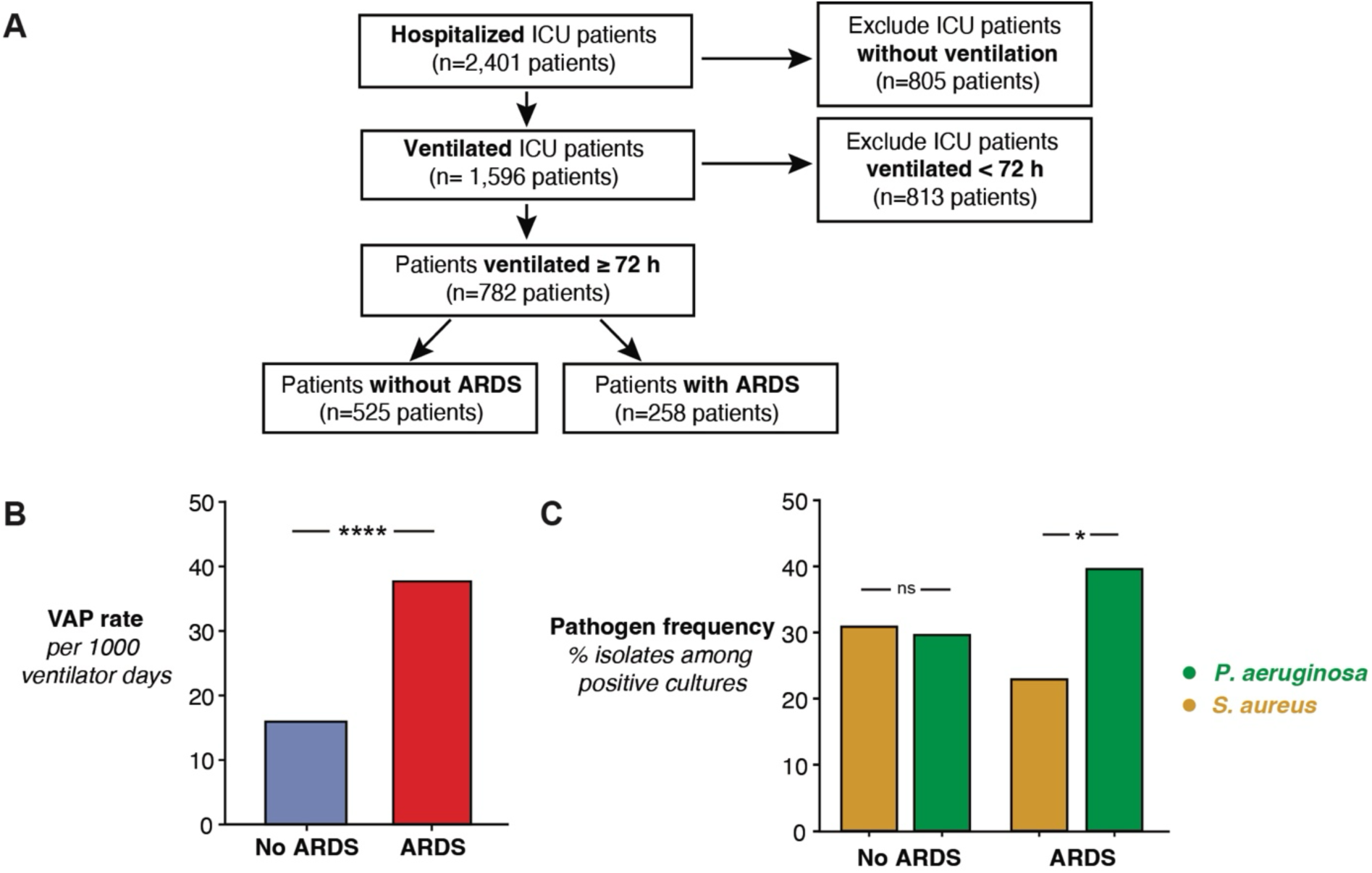
Incidence and most frequent etiological agents of ventilator-associated pneumonia in patients with and without ARDS, **a,** Inclusion criteria for a retrospective observational cohort of 783 mechanically ventilated patients. Criteria included admission to intensive care units within the University of Michigan Hospital System between 2010 and 2017 and receipt of ≥ 72 hours of invasive mechanical ventilation. ARDS status was clinically adjudicated using modified Berlin criteria, b, Incidence of ventilator-associated pneumonia among 783 mechanically ventilated patients who did or did not meet Berlin criteria for ARDS. Rates are normalized to 1000 ventilator days. Significance determined by two-sided independent t test. c, Percentage of positive cultures identified as Pseudomonas aeruginosa or Staphylococcus aureus, stratified by ARDS status. Difference between P. aeruginosa and S. aureus determined by one-sided independent t test. Significance key: ns p > 0.05; * p ≤ 0.05; ** p ≤ 0.01; *** p ≤ 0.001; **** p ≤ 0.0001.

### Airspace-derived soluble factors enhance P. aeruginosa growth in bronchoalveolar lavage fluid from mice with ALI

We then set out to determine if *P. aeruginosa* growth is enhanced by soluble, acellular factors in the injured lung microenvironment. To do this, we used an established murine model of oxygen-induced lung injury, which recapitulates the major biochemical and histological hallmarks of ALI,^26,27^ including alveolar leak, the onset of which occurs on day 3 and is detectable by quantification of protein and IgM in bronchoalveolar lavage fluid (BALF; **Fig. 2a,b**). After collecting BALF from mice in this model, we cultured *P. aeruginosa ex vivo* in acellular BALF and measured bacterial growth via quantitative plating and spectrophotometry. We found that *P. aeruginosa* growth was significantly enhanced in BALF from injured mice on day 3, the same timepoint as alveolar leak onset (**Fig. 2c,d**). From these experiments, we concluded that soluble, acellular factors promote *P. aeruginosa* growth in acutely injured murine lungs.

**Fig. 2:**
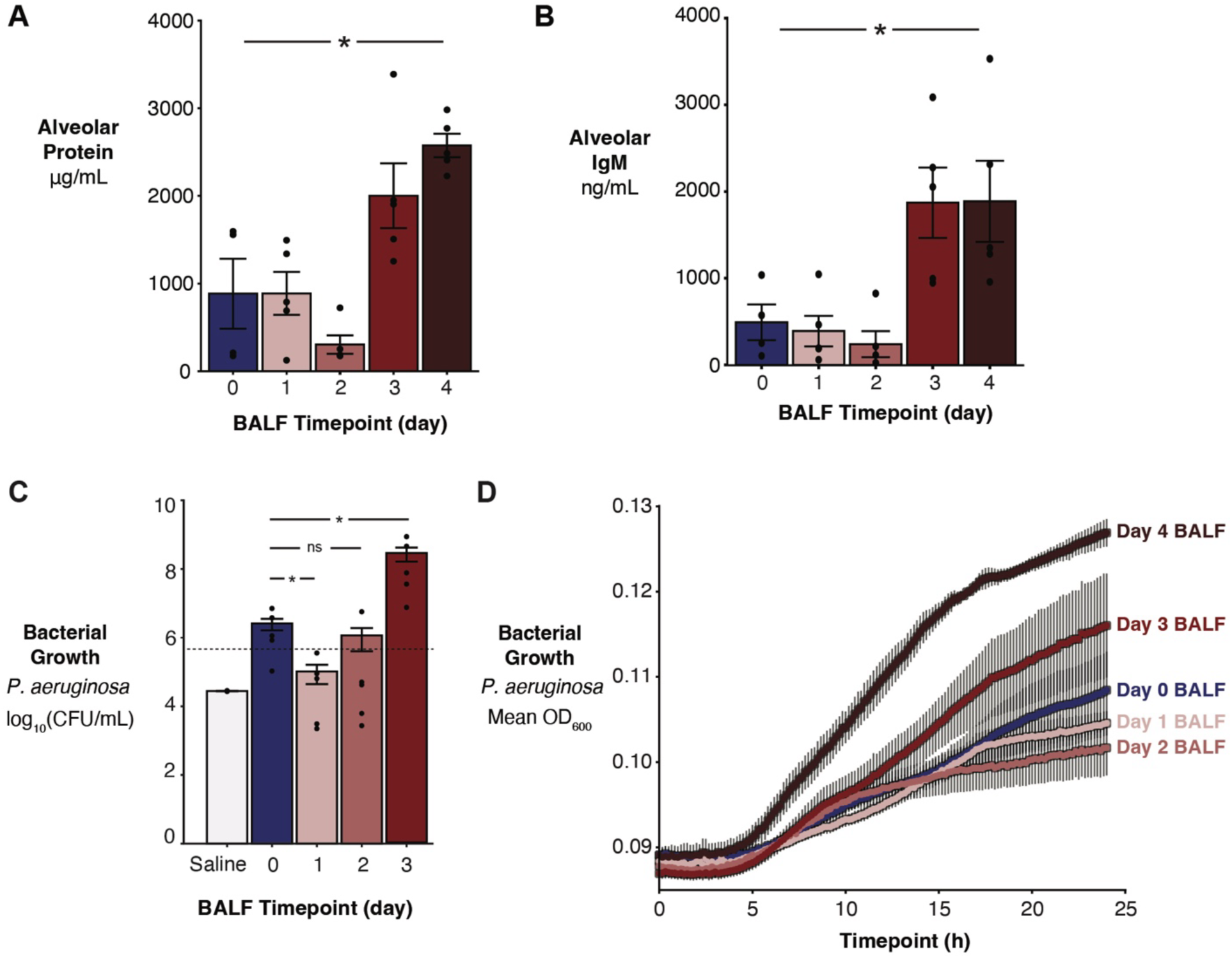
An ex vivo model of secondary bacterial pneumonia recapitulates increased growth of *P. aeruginosa* in BALF from mice with ALI. **a,b,** Alveolar leak in a murine model of oxygen-induced lung injury, as quantified by **(a)** total protein and **(b)** total IgM in BALF. Significance bar indicates the result of a Kruskal-Wallis rank sum test, c, Endpoint quantitative plating of an 8-hour *P. aeruginosa* culture in BALF from a murine model of oxygen-induced lung injury. Significance bars indicate the result of a pairwise Wilcoxon test, **d-f,** Spectrophotometric analysis as measured by optical density at 600 nm of *P. aeruginosa* over 24 hours in BALF from a murine model of oxygen-induced lung injury. Bacterial growth is indicated by optical density over time **(d)**, maximum optical density **(e)**, and doubling time **(f)** of *P. aeruginosa* cultured in murine BALF. Significance bars indicate the result of Kruskal-Wallis rank sum tests. Significance key: ns p > 0.05; * p ≤ 0.05; **p ≤ 0.01; *** p ≤ 0.001; **** p ≤ 0.0001.

### BALF from mice with ALI is enriched with an increased diversity and quantity of metabolites

Given that differential bacterial growth can be explained by nutrient limitation (modeled using serial dilutions of rich media; **Extended Data Fig. 1**), we hypothesized that soluble factors supporting *P. aeruginosa* growth in BALF from mice with ALI may be related to nutrient availability. To examine the nutrient content of BALF from mice with ALI, we compared metabolite profiles of murine BALF from healthy or injured (72 h hyperoxia exposure) mice using ^1^H nuclear magnetic resonance (NMR) spectroscopy (**Extended Data Fig. 2**). We found that the BALF metabolite profiles from injured mice were distinct from those of healthy mice by principal component analysis (**Fig. 3a**). BALF from injured mice contained higher concentrations of many metabolites; no detectable compounds were enriched in BALF from healthy mice (**Fig. 3b**). 3-hydroxybutyrate, a ketone body primarily produced by the liver and released into the blood during fatty acid oxidation, was the most relatively enriched compound in BALF from injured mice (**Fig. 3c**). Other BALF compound concentrations that were altered by ALI included glucose, lactate, and the branched-chained amino acids (BCAAs; **Fig. 3d-h**). Together, these results support the hypothesis that BALF from mice with ALI is enriched with metabolites that may serve as nutrient sources to support *P. aeruginosa* growth in the injured lung microenvironment.

**Fig. 3.**
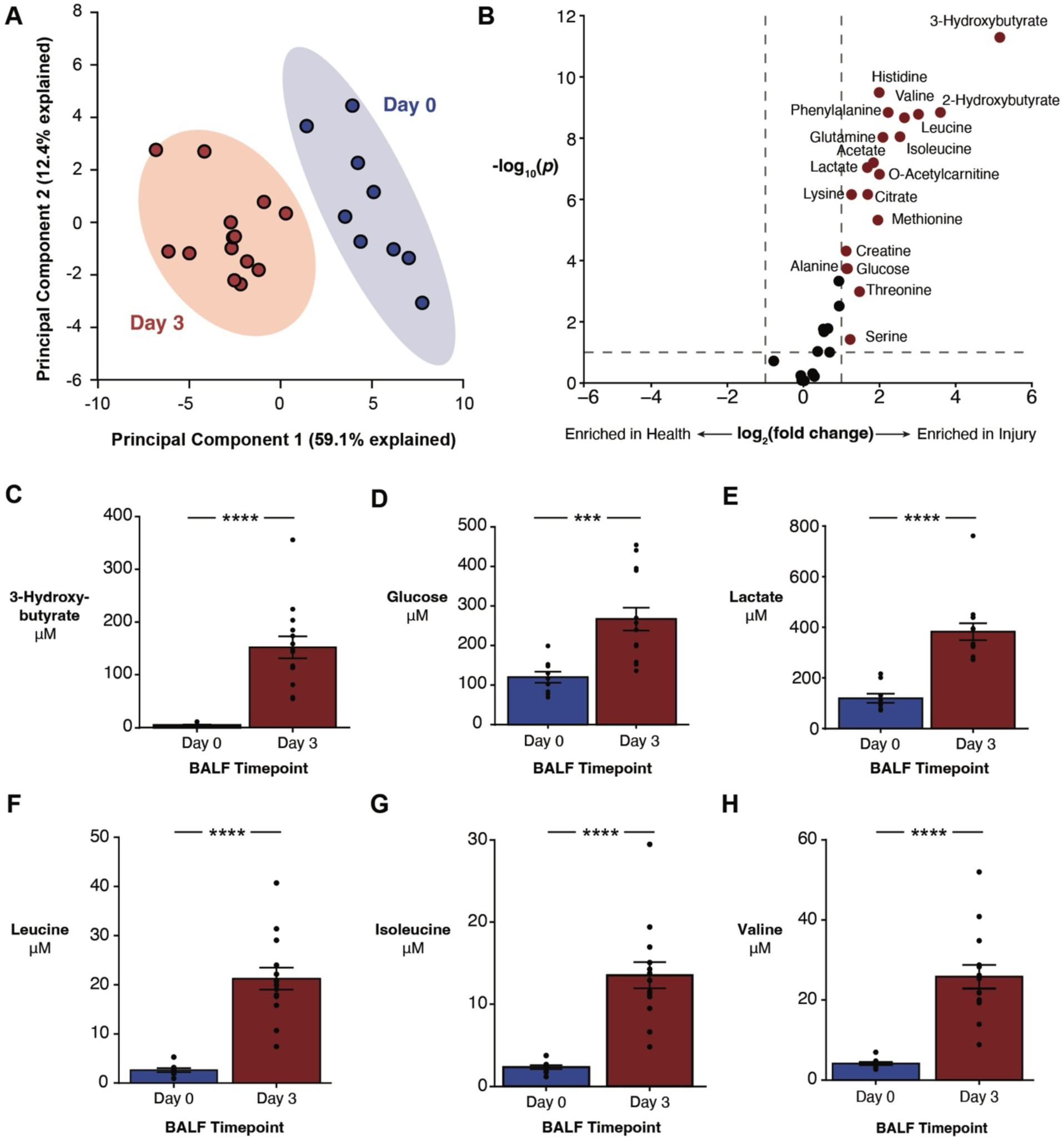
BALF metabolite profiles from mice with ALI are distinct from those in health and enriched with a variety of metabolites. **a,** Principal component analysis of murine BALF metabolite profiles characterized via NMR spectroscopy during health (day 0) and acute lung injury (day 3). Dots represent individual mice and ellipses represent 95% confidence interval, **b.** Volcano plot of murine BALF metabolites identified via NMR spectroscopy in injured lungs (day 3) compared to health (day 0). Dotted lines represent thresholds for fold-change (≥ 2-fold change in concentration, vertical) and significance (FDR-corrected p ≤ 0.1, horizontal). Significantly enriched features are indicated by red points and labeled with the metabolite name, **c-h**, Concentrations of significantly enriched metabolites identified in murine BALF during health (day 0) and acute lung injury (day 3), including 3-hydroxybutyrate **(c)**, glucose **(d)**, lactate **(e)**, and the branched chain amino acids leucine **(f),** isoleucine **(g),** and valine **(h).** Significance bars represent the results of a two-sided independent t test. Significance key: ns p > 0.05; * p ≤ 0.05; **ps ≤ 0.01; *** p ≤ 0.001; **** p ≤ 0.0001.

### Alveolar edema is a major source of metabolite enrichment in BALF from mice with ALI

Having found that BALF from mice with ALI promotes *P. aeruginosa* growth and is enriched with metabolites, we turned to the origin of BALF metabolite enrichment. We hypothesized that alveolar edema, derived from blood leaking into the airspace across a compromised blood-air barrier, is a major contributor to increased metabolite availability in injured lungs. To do this, we first considered the BALF metabolites of conventional and germ-free mice over time. As we have previously published,^26^ germ-free mice experienced delayed onset of oxygen-induced ALI, allowing us to compare if changes in BALF metabolite concentration correspond to the *timing* of alveolar leak (**Fig. 4a**). Consistent with alveolar leak timing, we found that the total BALF metabolite concentration in conventional mice increases from baseline starting at day 3, whereas BALF total metabolite concentration increases from baseline on day 4 in germ-free mice (**Fig. 4b**). Furthermore, the timing of BALF metabolite enrichment aligned with the timing of changes in BALF metabolite composition in both conditions (**Extended Fig. 3a**). We then examined if the *magnitude* of alveolar leak (cross-sectionally across mice) corresponds with metabolite enrichment during ALI by calculating correlation coefficients for each metabolite with paired IgM and protein concentrations. We found that most correlation coefficients were positive, regardless of the injury marker used (**Fig. 4c, Extended Data Fig. 3b**), indicating that alveolar leak directly correlates with alveolar metabolite concentration. Metabolites strongly correlated with protein and IgM BALF concentrations included glucose, lactate, 3-hydroxybutyrate, and the BCAAs (**Supplementary Data Table 1**). Together, these findings indicate that BALF metabolite enrichment during ALI occurs at alveolar leak onset and correlates with the severity of leak.

**Fig. 4:**
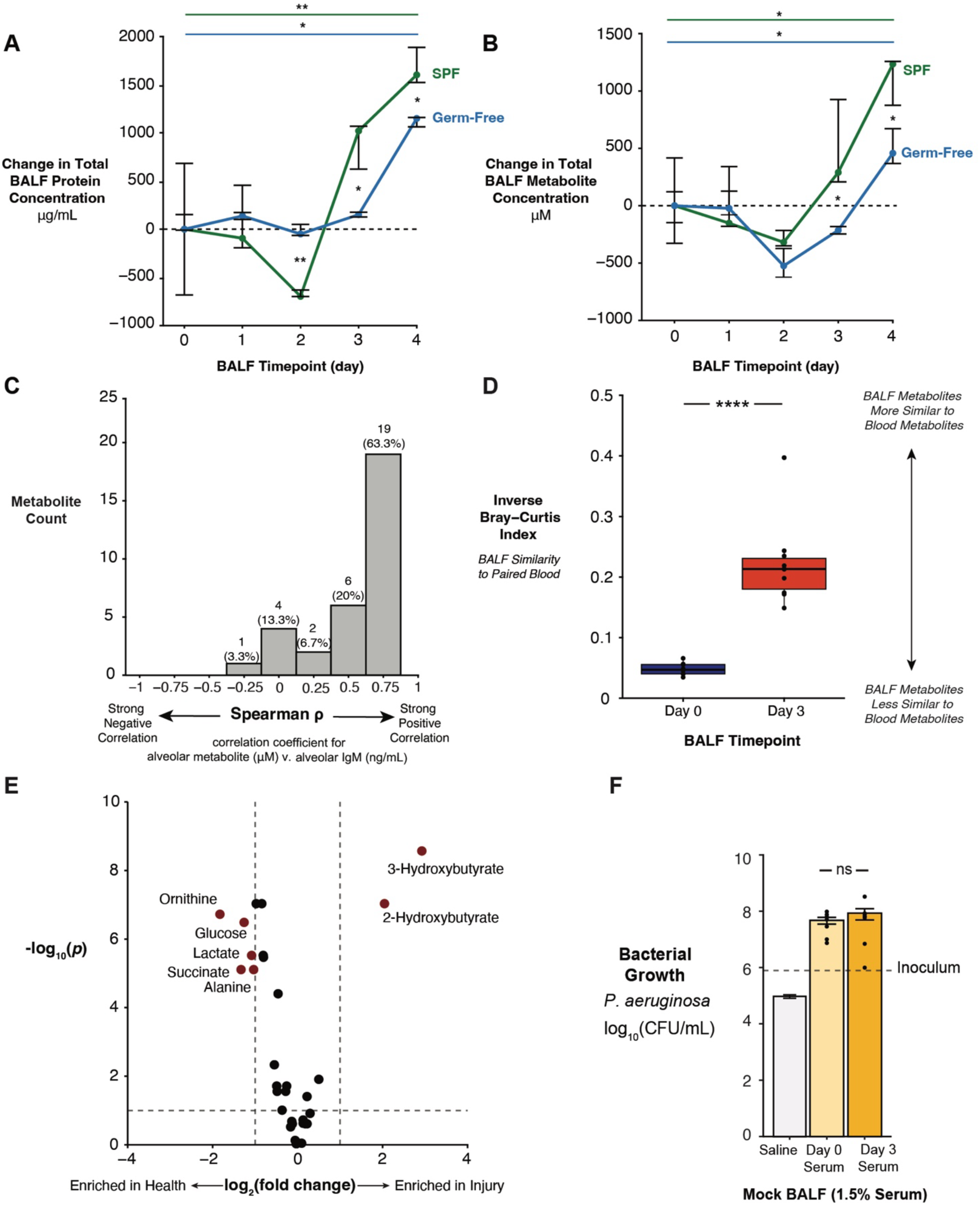
Pulmonary edema is a major source of metabolites enriched during alveolar leak. **a,b,** Change in alveolar leak, measured by total protein **(a)** and total metabolite concentration **(b)** during a 4-day timecourse of hyperoxia. Dots represent median and bars represent IQR. Across-group significance bars represent the results of Kruskal-Wallis rank sum tests, colored by microbiome group. Between-group significance stars represent the results of between-group Wilcoxon rank sum tests. **c,** Histogram of Spearman ρ coefficients for BALF lgM concentration with concentrations of each BALF metabolite detected via NMR. **d,** Inverse Bray-Curtis similarity index for metabolites detected via NMR in paired BALF and blood samples. Significance bar represents the results of a Wilcoxon rank sum test. **e,** Volcano plot of murine blood metabolites detected via NMR in injured mice (day 3) compared to health (day 0). Dotted lines represent thresholds for fold-change (≥ 2-fold change in concentration, vertical) and significance (FDR-corrected p ≤ 0.1, horizontal). Significantly enriched features are indicated by red points and labeled with the metabolite name. f, *P. aeruginosa* growth in mock BALF simulated by diluting serum in PBS (1.5% serum), measured by quantitative CFU plating, Significance bar represents the results of a Wilcoxon rank sum test. Significance key: ns p > 0.05; * p ≤ 0.05; ** p ≤ 0.01; *** p ≤ 0.001; **** p ≤ 0.0001.

Given this indirect evidence that pulmonary edema may be a source of metabolites in injured lungs, we then directly examined whether the blood is a source of these metabolites. First, we tested whether the BALF metabolite profiles of injured mice are similar to paired blood metabolite profiles using a similarity metric, the inverse Bray-Curtis index. At day 3 of hyperoxia in conventional mice, we found that BALF metabolites were more similar to metabolites in paired blood samples than for the same comparison in healthy mice (**Fig. 4d**). Increased similarity between BALF and blood metabolite profiles occurred only during alveolar leak in timecourse experiments (*i.e*., at 3 and 4 days in conventional mice and 4 days for germ-free mice; **Extended Data Fig. 3c**). Next, we examined how blood metabolite profiles change during onset of alveolar leak. Strikingly, 3-hydroxybutyrate was the most enriched in blood from mice with ALI relative to healthy mice (**Fig. 4e**). Even for metabolites enriched in blood from healthy mice (*e.g.*, glucose, lactate), the absolute concentrations of blood metabolites were consistent with the observed concentrations in BALF when accounting for the dilution that occurs during the lavage procedure (**Supplementary Data Table 2**).

Finally, to confirm that blood metabolites can support bacterial growth, we formulated “mock BALF” by diluting serum from healthy and injured mice with PBS so that metabolite concentrations in these “mock BALF” aliquots were similar to that of BALF from mice with ALI, and used these “mock BALF” aliquots as culture media for *P. aeruginosa*. Interestingly, we found no difference in bacterial growth between “mock BALF” formulated from serum of healthy and injured mice (**Fig. 4f**), indicating that the increased concentration of serum-derived metabolites in the injured lung is due to loss of barrier integrity, rather than availability of metabolites only present in injured serum, supports the growth of *P. aeruginosa* in the injured lung microenvironment. Together, these data support the hypothesis that airspace flooding with pulmonary edema containing blood-derived metabolites contributes to alveolar metabolite enrichment during ALI in our murine model.

### Edema-derived metabolites support P. aeruginosa growth in BALF from mice with ALI

Having established that increased *P. aeruginosa* growth in BALF from mice with ALI coincides with enrichment of blood-derived metabolites, we then examined *P. aeruginosa* gene expression to determine the metabolic pathways involved in enhanced growth of *P. aeruginosa* during secondary infection. To do this, we cultured *P. aeruginosa* in pooled BALF from multiple healthy or injured mice to mid-log phase and performed RNA sequencing. First, we tested whether *P. aeruginosa* gene expression changed in BALF from mice with ALI by performing differential gene expression analysis. We found that 1,812 genes were upregulated and 1,939 genes were downregulated when we cultured *P. aeruginosa* in BALF from mice with ALI compared to BALF from healthy mice (**Fig. 5a**). Top differentially expressed genes (DEGs) up in BALF from injured lungs included genes involved in toxin production and carnitine, lactate, and amino acid metabolism. In contrast, DEGs up in BALF from healthy lungs included genes involved in ethanol oxidation (**Supplementary Data Table 3**).

**Fig. 5.**
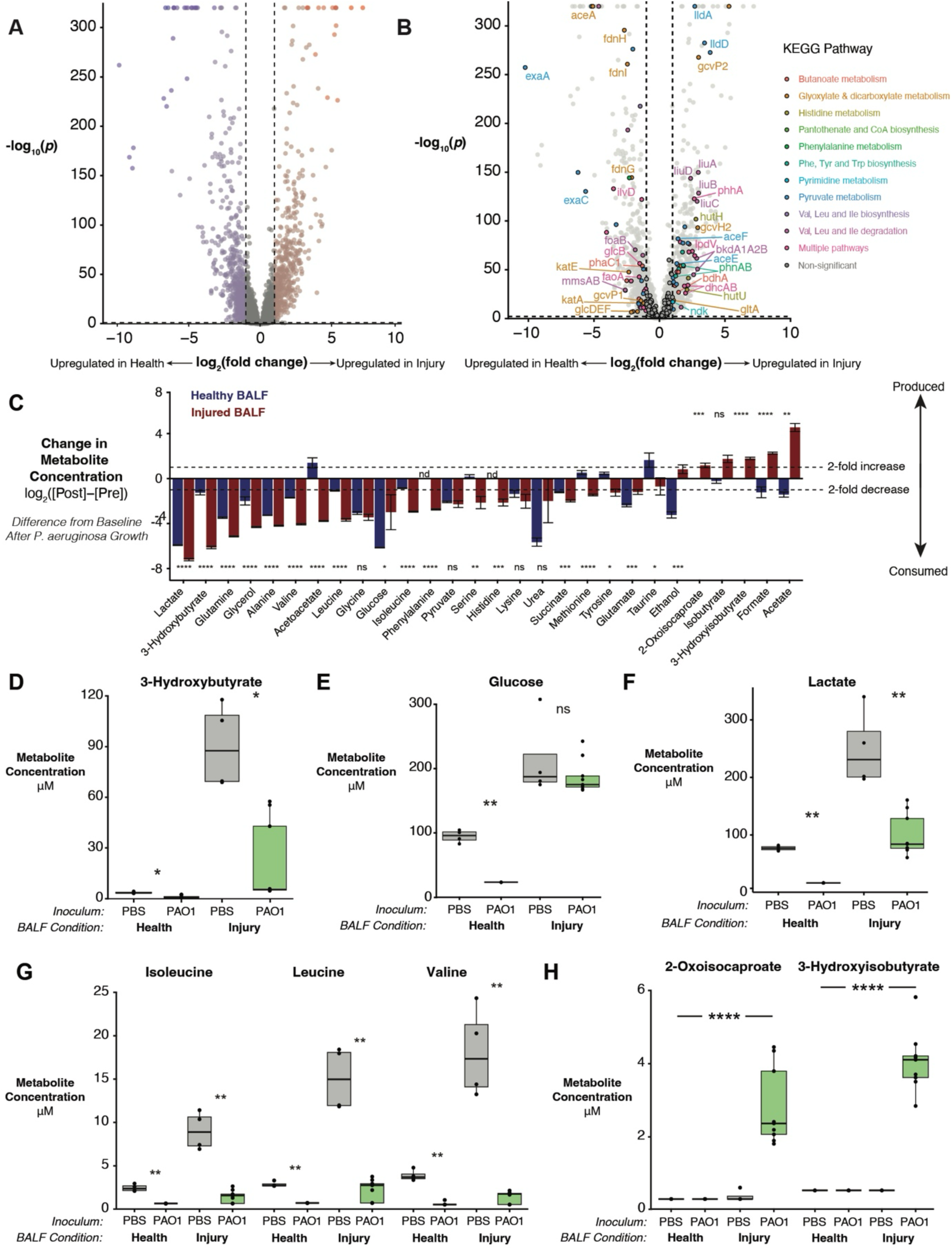
*P. aeruginosa* gene expression and metabolite uptake is distinct when cultured in BALF from mice with and without ALI. **a,b,** Volcano plot of genes differentially expressed by *P. aeruginosa* when cultured in BALF from mice with ALI relative to culture in BALF from healthy mice. Dotted lines represent thresholds for fold-change (≥ 2-fold change in expression, vertical) and significance (FDR-corrected p ≤ 0.01, horizontal). Genes that meet or exceed significance and fold-change thresholds are colored by injury status **(a)** or FDR-ranked top 10 KEGG pathway predicted by Metaboanalyst pathway analysis **(b). c,** Fold-change analysis of metabolites detected by NMR before and after *P. aeruginosa* culture in BALF from mice with (red) and without (blue) ALI. Bars are ordered by increasing value within the injured BALF group, bars represent mean fold-change, and error bars signify SEM. Statistical significance was determined via FDR-adjusted Student’s t test. **d-h,** Concentrations of metabolites identified in healthy and injured murine BALF after 10-h incubation with *P. aeruginosa* (PAO1) or vehicle (PBS), including 3-hydroxybutyrate **(d),** glucose **(e),** lactate **(f),** the branched chain amino acids **(g)**, and branched chain amino acid breakdown intermediates (h). Statistical significance was determined via pairwise Wilcoxon rank sum test with Benjamini-Hochberg correction for multiple comparisons **(d-f)** or Kruskal-Wallis rank sum test **(g,h)**. Significance key: ns nonsignificant; * p < 0.05; *** p < 0.001; **** p < 0.0001.

We next asked how many of the 3,751 DEGs were associated with metabolic pathways *and* differed between health and ALI (**Extended Data Fig. 4a**). To do this, we predicted which *P. aeruginosa* metabolic pathways should differ between conditions using pathway analysis of murine BALF metabolite profiles (*i.e.*, metabolite data shown in **Fig. 3a,b**), and then compared these results to actual changes in metabolic pathways observed in the transcriptomic data. Nearly one-third of metabolism-related gene expression changes in *P. aeruginosa* were predicted by differences in the BALF metabolite profiles between conditions (*i.e.*, 31.9% of DEGs were associated with KEGG pathways predicted to differ between conditions by metabolomic pathway analysis; **Supplementary Data Table 4**). DEGs associated with the top, statistically-ranked metabolic pathways in this analysis included genes involved in amino acid and butanoate degradation and assimilation of carbon compounds into central metabolism (**Fig. 5b**). Four metabolic pathways predicted by the injured BALF metabolite profiles were independently verified by overrepresentation analysis of KEGG pathways in the transcriptomic data (**Extended Data Fig. 4b**), demonstrating congruence between expected and observed metabolic responses of *P. aeruginosa* to the injured lung environment. Functional transcriptomic analysis of KEGG modules also identified differences in virulence factor expression between conditions, including increased expression of genes related to pyoverdine biosynthesis and the type 3 secretion system in injured BALF (**Extended Data Fig. 4c,d**). Together, these results indicate that changes in metabolism- and virulence-related gene expression by *P. aeruginosa* in the injured lung are at least partly attributable to the metabolic milieu of its environment.

Having identified transcriptome-level changes during culture of *P. aeruginosa* in BALF from mice with ALI, we then sought to confirm the activity of identified metabolic pathways at a functional level (*i.e.*, which metabolites are used by *P. aeruginosa* in injured BALF). To do this, we profiled metabolites before and after *P. aeruginosa* growth in BALF from healthy and injured mice, at the same mid-log-phase endpoint used in the transcriptomic analysis. We found that P. aeruginosa depletes many metabolites from BALF, with 23 of the 45 detectable metabolites in this analysis displaying at least a two-fold change in BALF from either condition (**Fig. 5c**). Notable examples include 3-hydroxybutyrate, which was only available in and depleted from BALF from injured mice (**Fig. 5d**), and glucose, which was only depleted by *P. aeruginosa* in BALF from healthy mice despite availability in both conditions (**Fig. 5e**). Several metabolites were depleted in BALF from both conditions (but to a greater extent in injury due to increased availability), including lactate and BCAAs (**Fig. 5f,g**). Furthermore, we detected five metabolites which were *produced* in at least one condition, including BCAA breakdown intermediates (**Fig. 5h**). Together, these results indicate that altered *P. aeruginosa* metabolite utilization – including increased use of metabolites available in both conditions and use of injury-specific metabolites – supports increased *P. aeruginosa* growth in injured lungs.

### Metabolite enrichment and enhanced P. aeruginosa growth coincides with alveolar leak in BALF from mechanically ventilated patients

Next, we investigated if the enhanced growth of *P. aeruginosa* in BALF from mice with ALI also applies to BALF from humans with ALI/ARDS. To do this, we cultured *P. aeruginosa* in BALF collected within 12 hours of intubation from a prospective cohort of mechanically ventilated ICU patients (n=113, **Fig. 6a, Extended Data Table 3)**. Similar to our murine findings, we hypothesized that *P. aeruginosa* growth would correspond with alveolar leak. We first grouped patients by P/F ratio at the time of BALF collection. P/F ratio, defined as the ratio of arterial oxygen partial pressure (PaO_2_) to fractional inspired oxygen (FiO_2_), is a nonspecific index of impaired oxygenation, a criterion in the Berlin definition of ARDS, and the most common index of ARDS severity.^25^ We found that BALF from mechanically ventilated patients with a P/F ratio ≤ 300 (*i.e.*, patients with possible ALI/ARDS) increased *P. aeruginosa* growth more than BALF from patients with a P/F ratio above 300 (*i.e.*, patients without ALI/ARDS), as quantified by maximum optical density (OD) in a 24-hour culture (**Fig. 6b**). The correlation between P/F ratio and maximum OD was negative but weak (**Fig. 6c**, Spearman ρ = −0.11, *p* = 0.58), reflecting both the limitations of the P/F ratio (which is nonspecific and lowered by processes other than alveolar leak) and the wide range of OD values within the binary bins. To more directly examine the relationship between alveolar leak and bacterial growth, we quantified BALF IgM levels and found a strong positive correlation between BALF IgM concentration and maximum OD (**Fig. 6d**, Spearman ρ = 0.63, *p* = 7.7 x 10^-14^). Together, these data indicate the relationship between alveolar leak and *P. aeruginosa* growth in BALF is also evident in mechanically ventilated patients.

**Fig. 6.**
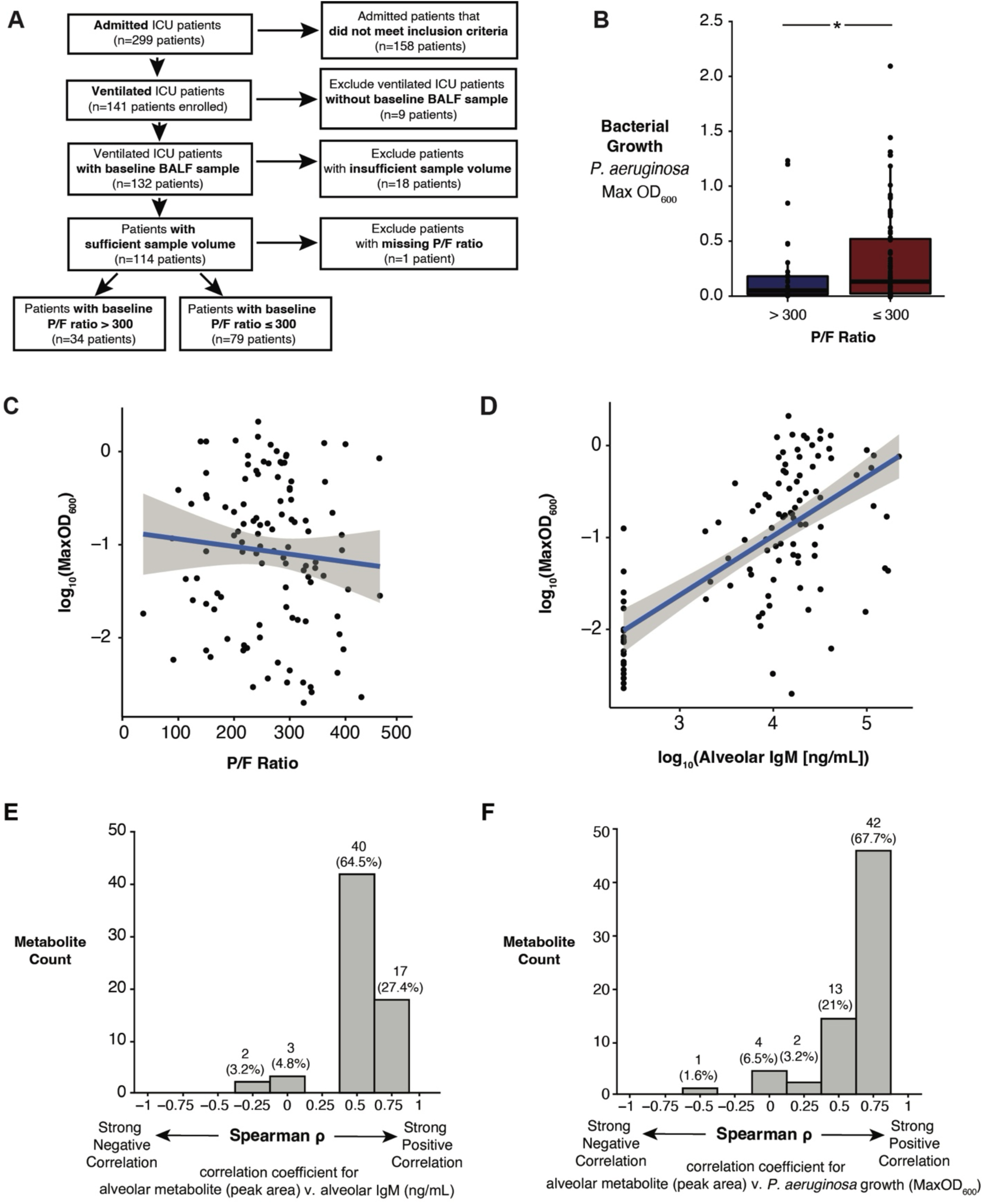
*P. aeruginosa* growth in BALF from mechanically ventilated patients positively correlates with alveolar leak and metabolite availability. **a,** Inclusion criteria for a prospective human cohort of 114 mechanically ventilated patients. Criteria included admission to intensive care units Clfnica Universidad de La Sabana from 2020 and 2022 and receipt of ≥ 12 hours of mechanical ventilation. Baseline P/F ratio was calculated from electronic medical record data at the time of BALF collection. **b,** Maximum optical density of *P. aeruginosa* cultured in human BALF aliquots from patients with baseline P/F ratio >300 or ≤ 300. Significance bar represents the result of a two-sided independent I test. **c,d,** Correlation of maximum optical density of *P. aeruginosa* cultured in human BALF aliquots with P/F ratio (c) and BALF lgM concentration (**d**). Blue line with gray halo represents linear model with 95% confidence interval. **e,f,** Histogram of Spearman p coefficients for correlation of concentrations of each BALF metabolite detected via GC-QTOF-MS with human BALF lgM concentration **(e)** and maximum optical density of *P. aeruginosa* cultured in paired human BALF aliquots (f). Significance key: ns p > 0.05; * p ≤ 0.05; ** p ≤ 0.01; *** p ≤ 0.001; **** p ≤ 0.0001.

Finally, we determined which metabolites are present in BALF from mechanically ventilated patients and their relationship with alveolar leak and *ex vivo* growth of *P. aeruginosa*. To do this, we measured metabolites in patient BALF using an untargeted gas chromatography-mass spectrometry-based metabolomics approach, which identified 62 metabolites (**Extended Data Fig. 5a**). To compare BALF metabolites with severity of alveolar leak, we calculated the Spearman correlation coefficient of detected metabolites with alveolar IgM and found a positive correlation for nearly all identified metabolites (**Fig. 6d**). We also calculated the Spearman correlation coefficient of metabolites with *P. aeruginosa* maximum OD and found a similar pattern of strong positive correlation of most metabolites with bacterial growth (**Fig. 6e**). Notably, 3-hydroxybutyrate and the BCAAs (metabolites that positively correlated with alveolar leak and are used by *P. aeruginosa* in BALF from mice with ALI) had strong positive correlations with *P. aeruginosa* growth in this human dataset (**Extended Data Fig. 5b-e).** These findings further corroborate that alveolar leak is a major source of metabolites that can fuel *P. aeruginosa* growth in acutely injured lungs.

## Discussion

In this study, we found that in both humans and mice, ALI/ARDS increases nutrient availability in the alveolar microenvironment, contributing to *P. aeruginosa* growth in the acutely injured lung. In experimental lung injury and humans with respiratory failure, changes in BALF metabolites and bacterial growth are attributable to severity of alveolar leak, indicating that serum-derived edema is a major nutrient source for respiratory pathogens in ALI/ARDS. Our findings establish that the alveolar metabolic microenvironment is an important ecological component of pneumonia pathogenesis in ALI/ARDS and identify potential targets for therapeutic interventions for pneumonia in this patient population.

Our findings expand the conceptual understanding of secondary pneumonia pathogenesis beyond its traditional pathophysiological framework. Whereas prior studies have focused on two factors contributing to the increased susceptibility of acutely injured lungs to secondary bacterial infection – pathogen virulence and host immunity – our study reveals that *ecological* factors contribute to pneumonia pathogenesis as well. Studies from the 19th century on have affirmed the role of bacterial virulence in facilitating pneumonia, with molecular studies identifying genetically-encoded virulence factors (*e.g.*, exotoxin A in *P. aeruginosa*) that contribute to successful infection and select species repeatedly identified as causative agents of secondary lung infections (*e.g.*, *P. aeruginosa*, *S. aureus*, *Klebsiella pneumoniae*, *Escherichia coli)*. Impaired immune defenses also play a role in establishing secondary infection. Healthy lungs contain a multi-layered arsenal of immunological mechanisms that maintain the near sterility of the airspace, including anatomical (*e.g.*, mucociliary escalator, cough), molecular (*e.g.*, antimicrobial peptides, Toll-like receptors) and cellular (*e.g.*, alveolar macrophages, neutrophils) defenses. While these immunological protections maintain the lung’s functionality by preventing the physical obstruction of gas exchange, dysfunction of these defenses permits the accumulation of microbial biomass, a critical step in establishing infection. Recent research in pulmonary microbial ecology^28^ has suggested that a third factor – altered rates of bacterial reproduction due to changes in the lung microenvironment – contributes to lung microbiome dysbiosis (of which secondary pneumonia is an extreme example).^24^ Our study provides proof-of-concept evidence that changes in the alveolar metabolic microenvironment caused by leak of serum-derived edema into the airspace facilitate increased growth of bacteria that frequently cause secondary pneumonia.

Our findings suggest that bacterial fitness for the ALI/ARDS-associated metabolic microenvironment may explain why such a small subset of species causes most secondary infections. Our retrospective cohort of mechanically ventilated ICU patients found that *P. aeruginosa* was the most frequent cause of VAP in patients with ARDS, consistent with other reports.^3,29^ *P. aeruginosa’*s prevalence as a cause of secondary pneumonia in patients with ALI/ARDS is consistent with available metabolites we detected in specimens from injured lungs. *P. aeruginosa* prefers to metabolize nonfermentable carbon sources, which can be completely oxidized with minimal use of resources (*e.g.*, lactate, acetate) over glucose,^30,31^ a phenomenon termed “reverse diauxie” in contrast to *E. coli’s* preferential utilization of glucose over other substrates.^32^ More generally, pseudomonads are known for preferential metabolism of organic acids and amino acids over carbohydrates, a property observed for *P. aeruginosa* in the context of planktonic and biofilm growth,^33^ *ex vivo* growth in burn wound exudate^34^ and cystic fibrosis sputum,^35^ during cross-feeding with the commensal *Rothia mucilaginosa,*^36^ and now in *ex vivo* culture with BALF from patients and mice with ALI/ARDS.

Though biochemical and metabolomic studies of BALF from patients with ALI/ARDS are few, studies to date have suggested a global increase in metabolite availability, an effect we observed in our murine studies.^37,38^ Compared to healthy volunteers and patients with non-ALI/ARDS respiratory failure, patients with ALI/ARDS have increased concentrations of BALF protein,^39,40^ glucose,^41^ lactate,^42^ and amino acids,^40^ which we also identified in murine BALF. While some evidence to date links changes in the pulmonary metabolic microenvironment to pneumonia risk (*e.g.*, bronchial aspirate glucose concentration and staphylococcal infection,^41^ influenza-induced sialylated mucin availability and streptococcal infection^14^), our study fills a key knowledge gap by focusing on *P. aeruginosa*’s response to nutrient availability during ALI/ARDS.

A notable theme in our data was related to 3-hydroxybutyrate, a ketone body primarily produced in the liver during host nutritional stress. In our experimental model of ALI/ARDS, 3-hydroxybutyrate is present in the blood during alveolar leak, suggesting its high concentration and dramatic fold-change difference in BALF from injured mice is largely driven by serum-derived edema. We also detected 3-hydroxybutyrate in BALF from mechanically ventilated patients, suggesting translatability of this murine finding to human disease. Pseudomonads, including *P. aeruginosa*, import 3-hydroxybutyrate via a putative H^+^ gluconate symporter family transporter (PA2004), dehydrogenate 3-hydroxybutyrate via BdhA, and shunt the resulting acetoacetate into central metabolism (KEGG pathway pae00650), a process regulated in *P. aeruginosa* by the enhancer-binding protein HcbR and alternative sigma factor RpoN.^43^ In our analyses, *bdhA* expression increased and 3-hydroxybutyrate was depleted when *P. aeruginosa* was grown in BALF from mice with ALI, suggesting that 3-hydroxybutyrate may play an important role in acute infection during ALI/ARDS. Recently, Tomlinson and colleagues published that 3-hydroxybutyrate availability in the airways plays a role in establishing *P. aeruginosa* tolerance during chronic infection.^44^ Together, these results suggest a dual role for 3-hydroxybutyrate across distinct infection contexts and emphasize the importance of considering systemic metabolites in pneumonia.

Our study has several limitations. First, we used a laboratory *P. aeruginosa* strain as a model species for proof-of-concept; studies of other pneumonia-causing bacteria will be necessary to determine applicability across species. We have observed that *S. aureus* growth increases when cultured in BALF from hyperoxia-exposed mice relative to healthy controls, suggesting that our findings are not unique to *P. aeruginosa*.^26^ Second, *ex vivo* culture of *P. aeruginosa* in BALF does not recapitulate all aspects of the *in vivo* alveolar microenvironment. Because BALF is derived from diluting airspace contents, BALF metabolite concentrations are considerably lower than physiological concentrations in biofluids like airway surface liquid or pulmonary exudate and, as such, are not directly comparable to source samples (*e.g.*, blood). Though future *in vitro* and *in vivo* experiments are warranted, our data are consistent with reports of increased bacterial burden and mortality in hyperoxia-exposed mice infected with *P. aeruginosa.*^12,45^ Third, we used NMR spectroscopy for metabolite profiling of murine BALF and blood. NMR only detects metabolites at micromolar-scale concentrations (*i.e.*, less sensitive than mass spectrometry-based platforms). Though our metabolomics strategy does not detect metals, hydrophobic compounds, or macromolecules – which also may serve as bacterial nutrients in injured lungs – we identified highly abundant metabolites that serve as major nutrient sources, providing a foundation for future study.

## Methods

### Ethics statement

All clinical investigations were conducted according to the principles expressed in the Declaration of Helsinki.

The institutional review board of the University of Michigan Healthcare System (HUM00104714) approved the retrospective observational human study protocol. The institutional review board examined the protocols for this de-identified analysis of patient data and assessed there to be no more than minimal risk with no need for a dedicated informed consent process.

The institutional review board of Clínica Universidad de La Sabana approved the prospective human study protocol (20190903, 468). All patients provided written informed consent. All methods and research procedures were performed in accordance with the local and international regulations for good clinical practices in clinical research and did not change the clinical treatment of the patients participating in the study as per local clinical guidelines. Clinical data for enrolled patients was gathered in REDCap and hosted securely on the Universidad de La Sabana server.

The University Committee on the Care and Use of Animals at the University of Michigan (PRO00007791, PRO00009673, PRO00011460) approved the animal studies in this manuscript. Laboratory animal care policies at the University of Michigan follow the Public Health Service Policy on Humane Care and Use of Laboratory Animals. Animals were assessed twice daily for physical condition and behavior. Animals assessed as moribund were humanely euthanized by CO_2_ asphyxiation.

### Retrospective human cohort analysis

We performed a retrospective cohort study of patients at the University of Michigan Hospital who received mechanical ventilation for at least 72 hours. The observational human cohort was identified via abstraction from the University of Michigan’s electronic medical record. Inclusion criteria were 1) age ≥ 18 years, 2) admission to the University of Michigan’s Critical Care Medical Unit, Surgical Critical Care Unit, Cardiac Intensive Care Unit, Burn Intensive Care Unit, Medical Surgical Moderate Care Unit, or Cardiovascular Moderate Care Unit between January 1, 2016, and December 31, 2017, and 3) received invasive mechanical ventilation for at least 72 hours. VAP, our primary outcome of interest, occurs after 48 hours of mechanical ventilation, and thus, the 72-hour time frame was chosen to identify a population with a non-zero probability of developing VAP.

Diagnosis of the acute respiratory distress syndrome (ARDS) was made using the Berlin Criteria^25^ and adjudicated by three critical care-trained physicians as previously described.^46^ VAP was diagnosed using the following streamlined version of the CDC surveillance criteria:^47^ 1) moderate or numerous leukocytes present in purulent sputum production after 48 hours of mechanical ventilation, 2) sustained increase in oxygen requirement (at least one of the following: 2 days of increasing daily minimum FiO2 ≥ 0.15 or 2 days of increasing daily minimum PEEP > 2.5 mm H_2_O) within a 3-day window of the respiratory culture collecting day (including one day before and one day after), 3) evidence of systemic inflammation present (at least one: fever [temperature > 38 °C] or hypothermia [temperature < 35°C) or leukocytosis [white blood cell count > 12,000], or leukopenia [white blood cell count < 4,000], and 4) pathogen growth on respiratory culture.

Lower respiratory specimens were collected via lavage (either bronchoalveolar lavage or non-bronchoscopic bronchoalveolar lavage [mini-BAL]) and cultured according to previously described clinical microbiology laboratory protocols.^26,48^ Briefly, specimens were plated and incubated for 72 hours on chocolate, sheep blood, and MacConkey agar. Colonies were identified and species reported if greater than 10^4^ CFU/mL grew or if less than 10^4^ CFU/mL grew but colonies were a single Gram-negative bacillus and the only reportable pathogen.

### Murine modeling

Eight-week-old female C57Bl/6 conventional, specific pathogen-free mice (n = 125) were purchased from the Jackson Laboratory (Bar Harbor, ME) and housed under specific pathogen-free conditions. Mice were allowed to acclimate for about a week before experiment start and harvested at 9 weeks of age. Mice were housed in a common animal housing room and did not receive independent ventilation during acclimation and experimentation.

Germ-free mice (n = 66) between the ages of 8 to 12 weeks were obtained via the University of Michigan Germ-Free Animal Core. Until the time of hyperoxia exposure, mice were housed in soft-sided plastic isolators to prevent exposure to microbes. Germ-free status was regularly confirmed via culture- and PCR-based monitoring of fecal and necropsy samples. Germ-free mice were placed in the hyperoxia chamber using dedicated microisolator cages with sufficient food and water for the duration of exposure. Thus, microisolator cages were kept sealed and sterile until the time of tissue harvest.

Hyperoxia was administered to mice by placing their cages in a sealed chamber (BioSpherix) with medical-grade 100% oxygen (0.1 to 99.9 ± 0.1%) delivered continuously via a ProOx 110 oxygen controller to maintain chamber oxygen levels. Adult mice were administered FiO2 95% for durations ranging from 24 to 96 hours.

### Murine sample collection and processing

Mice were sacrificed via CO_2_ asphyxiation, and diaphragm puncture was performed as a secondary euthanasia method. All tissue samples were collected using sterile technique; instruments were rinsed with ethanol and flamed between each sample type. BALF collected for metabolomics analysis was collected without using ethanol to spray down the fur before dissection (PBS was used instead), as samples collected with ethanol were found to display large ethanol peaks on NMR spectra and obscure peaks from other metabolites of interest.

Whole blood was collected via cardiac puncture; the right ventricle was punctured with a heparinized, sterile syringe attached to a 25G needle, and blood was drawn slowly to prevent ventricular collapse. Following blood collection, the blood was placed in a cryovial with 10 μl of 5000 U/mL heparin, rolled gently to mix, and flash frozen in liquid N_2_.

BALF was collected by dissection to expose the trachea, insertion of sterile tubing and connected needle and syringe, tightening of surgical thread around the intubated trachea to seal, and two rounds of instillation and aspiration of 1 mL of sterile phosphate-buffered saline. Each tubing-needle-syringe setup was rinsed thoroughly with sterile PBS between sample collections. Both serial lavages were pooled, yielding up to 2 mL BALF per mouse. Pooled BALF was centrifuged (16,500 x *g* for 30 min at 4°C) and the acellular supernatant was transferred to cryovials, flash frozen with liquid N_2_, and stored at −80°C until use. If samples were to be used for multiple purposes, pooled BALF was aliquoted into multiple vials to avoid multiple freeze-thaw cycles. Syringe rinses pre- and post-lavage and sterile PBS were collected, processed identically to BALF, and were used as negative controls for *ex vivo* culture and metabolomics experiments.

### In vitro and ex vivo bacterial culture

Murine or human BALF, sterilized via centrifugation, was divided into 100-μL aliquots in a 96-well plate and inoculated in duplicate with ∼10^4^ CFU of *Pseudomonas aeruginosa* strain PAO1. Uninoculated aliquots were used as negative controls to confirm the sterility of the fluid, with no detectable CFUs or increase in optical density, depending on the assay readout. For quantitative plating studies, each aliquot was incubated at 37°C with shaking at 125 rpm under normoxic conditions, plated in duplicate onto LB agar plates, and CFU were counted after 24 h. For spectrophotometric studies, prepared 96-well plates were placed in a temperature-controlled plate reader (cat. no. 30190085, Tecan US Inc., Morrisville, NC) under normoxic conditions, shaking at 140 rpm and incubated at 37°C for 24 h, with absorbance readings at 600 nm collected every 10 minutes.

The mock BALF experiment was conducted according to the same protocol for murine and human BALF except for the use of murine serum samples diluted to approximate the concentrations of IgM present in BALF during alveolar leak (1.5% serum in PBS). Mock BALF was created as follows: blood was collected from control and hyperoxia-exposed mice via retroorbital bleed into serum separator tubes (BD Microtainer® SST^TM^, cat. no. 365967) and processed according to manufacturer’s instructions. Undiluted serum samples were stored at −80°C until use and diluted upon thawing. Diluted serum was divided into 100-μL aliquots and used as growth media for *ex vivo* culture as described above for murine and human BALF.

To yield sufficient material for transcriptomic and metabolomic studies of *ex vivo* bacterial cultures, acellular BALF from multiple mice within the same treatment arm was pooled, and the microplate assay described above was scaled up to 10x culture volume. Pooled murine BALF was divided into 1000-μL aliquots in 5 mL sterile culture tubes and inoculated with ∼10^5^ CFU of *P. aeruginosa* str. PAO1. Uninoculated aliquots were used as negative controls to confirm BALF sterility. To confirm bacterial growth, a portion of each culture was then plated in duplicate onto LB agar plates after 10 hours of incubation at 37°C shaking at 125 rpm under normoxic conditions, and CFU were counted after 24 hours of subsequent growth. The remaining culture volume was transferred to a 2 mL microcentrifuge tube, centrifuged (17,110 x *g* for 5 min at RT), and the supernatant was removed and stored at −80°C until metabolomics processing. Two volumes of Qiagen RNA-protect (cat. no. 76506) was then added to the remaining bacterial cell pellet, vortexed vigorously, incubated at room temperature for five minutes, and then stored at −80°C until RNA isolation.

### Lung injury and inflammation

For murine studies, lung injury was assessed by quantification of total alveolar protein and alveolar IgM in BALF. Total alveolar protein was quantified colorimetrically via Bradford assay (Bio-Rad, cat. no. 5000006) with bovine serum albumin to generate the standard curve. Alveolar IgM was quantified using the IgM Mouse Uncoated enzyme-linked immunosorbent assay (ELISA) Kit (Thermo Fisher, cat. no. 88-50470-88). For human studies, lung injury was assessed by quantification of alveolar IgM in human BALF using the IgM Human Uncoated ELISA Kit (Thermo Fisher, cat. no. 88-50620-88).

### Murine BALF and whole blood NMR metabolomics

Acellular BALF was thawed at room temperature and ultrafiltered to remove macromolecules. Briefly, 500 μl of BALF was centrifuged (2,000 x *g* for 10 minutes at 4°C) to precipitate any sediment, and the resulting supernatant was transferred to Nanosep 3K centrifugal devices with an Omega filter membrane (Fisher Scientific, cat. no. 0D003C34) and centrifuged (14,000 x *g* for 20 min at 4°C). Any filtrate from the bottom of the centrifugal device was then transferred to a respectively labeled cryovial. Phosphate buffer in deuterium oxide (D_2_O; 50 μl) was then added to the filter membrane, vortexed for 30 seconds, and the sample was centrifuged again (14,000 x *g* for 25 min at 4°C). Any filtrate at the bottom of the centrifugal device was transferred to the same respective cryovial as the filtrate from the first spin. The post-filtration sample volume was measured using a 1mL glass serological pipet, phosphate (50 mM) buffer in D_2_O was added to an end volume of 500 μl, and sodium trimethylsilylpropanesulfonate (DSS-d6; 10 µL; IS-2, Chenomx, Edmonton, Alberta, Canada) was added as an internal standard. Samples were stored at −80°C until spectral acquisition.

Sodium heparin-preserved whole blood was thawed in an ice bath, and macromolecules were removed via 1:1 methanol/chloroform extraction as previously described,^49^ which precipitates proteins and separates hydrophilic and lipophilic fractions, the former of which was assayed in this study. Briefly, two volumetric equivalents of 1:1 methanol/chloroform solution were added to thawed whole blood, vortexed (30 s) to mix, sonicated (2 min), and incubated (−20°C for 20 min). Samples were then centrifuged (13,400 x *g* for 30 min at 4°C). Following centrifugation, aqueous supernatants were transferred to a new tube, frozen in liquid N_2_, and lyophilized (Labconco, Kansas City, MO) for at least 20 hours. Lyophilized samples were then resuspended in phosphate buffer (50 mM) in D_2_O (500 μL). After resuspension, sample volumes were measured, and phosphate buffer in D_2_O was added as needed to achieve a final volume of 500 μL for each sample. Finally, 4.99 mM DSS internal standard (50 μL) was added, and samples were stored at −80°C until spectral acquisition.

The resulting processed BALF and whole blood samples were assayed using ^1^H nuclear magnetic resonance spectroscopy. Spectra were acquired at the University of Michigan’s Biochemical NMR Core Laboratory on a Varian (now Agilent, Inc., Santa Clara, CA) 500MHz NMR spectrometer with a VNMRS console operated by host software VNMRJ 4.0. Spectra were recorded using 32 scans for whole blood or 256 scans for BALF of a proton-proton-NOESY pulse sequence, commonly called a METNOESY pulse sequence.^50,51^ Spectra were acquired at room temperature (295.45+/-0.3K) using a 5-mm Agilent “One-probe”. The pulse sequence is as follows: A 1s recovery delay, which includes a 990 ms saturation pulse of 80Hz induced field strength empirically centered on the water resonance, 2 calibrated 90 degree pulses, a mixing time of 100 ms, a final 90 degree pulse, and an acquisition period of 4s. Optimal excitation pulse widths were obtained by using an array of pulse lengths as previously described.^52^ DSS was used as the internal standard for whole blood and BALF.

Chenomx NMR Suite 8.6 software (Edmonton, Alberta, Canada) was used for spectral processing and metabolite quantification. Using the Chenomx Processor module, spectra were processed via phase shift adjustment, water region excision, baseline adjustment, and internal standard calibration. Metabolites were then identified and quantified using the Profiler module, which accounts for pH and integrates peaks using the known concentration of the DSS internal standard as a reference. Compounds are named using the 338-compound Chenomx v11 library or, if not identified in Chenomx, the 657-compound Human Metabolome Database v. 5.0. Compound concentrations were then volume-corrected to determine concentration in the original sample. In BALF, metabolites were removed from the dataset if they had ≥30% missingness in both healthy and injured groups. If a metabolite had ≤30% missingness in an injured group, it was retained. In blood, metabolites were removed from the dataset if they had ≥30% missingness, if they were known exogenous contaminants (*e.g.*, methanol), or if their concentrations are known to be affected by lyophilization (*e.g.*, acetate). Missing values were imputed using half the minimum detectable concentration of the compound in question. Imputation was performed separately for BALF and blood.

After these data reduction steps, the final set of identified metabolites in this series of experiments included 30 to 33 metabolites in BALF, 39 metabolites in whole blood, and 45 metabolites in BALF supernatant from *P. aeruginosa ex vivo* culture experiments. The final murine BALF and blood metabolomics datasets were analyzed using R version 4.4.0^53^ and Metaboanalyst 5.0.^54^ Multi-metabolite comparisons in Metaboanalyst were conducted on the log-transformed and autoscaled datasets.

### Bacterial RNA sequencing

Total RNA was isolated from bacterial cell pellets cultured *ex vivo* in acellular murine BALF using the Qiagen RNeasy kit (cat. no. 74104) according to manufacturer’s instructions, with optional enzymatic lysis via lysozyme digestion (Thermo Scientific cat. no. 89833, 15 mg/mL in TE buffer, incubated for 15 minutes at 37°C) and DNase I digestion (Qiagen cat. no. 79254).

Purified RNA was eluted in RNase-free water and stored at −80°C until downstream quality checks and sequencing. Purified RNA was submitted to the University of Michigan Advanced Genomics Core for quality checks, ribo-depletion, and bulk RNA sequencing using the Illumina NovaSeq platform. Quality of RNA samples was determined via Qubit RNA broad-range assay (Thermo Fisher cat. no. Q10211) and TapeStation (Agilent cat. no. 5067-5576). Samples were prepared for sequencing using the NEBNext rRNA Depletion Kit for bacteria (cat. no. E7850X), NEBNext UltraExpress RNA Library Prep Kit for Illumina (cat. no. E3330L), and NEBNext Multiplex Oligos for Illumina Unique dual kit (cat. no. E6448S). Briefly, 144 ng of total RNA was ribo-depleted, fragmented for 8 minutes as determined by RNA integrity number (RIN), and copied into first strand cDNA using reverse transcriptase and random primers. After cDNA synthesis, cDNA 3’ ends were adenylated, adapters were ligated, and products were purified and enriched by PCR to create the final cDNA library. Final libraries were checked for quantity and quality by Qubit hsDNA (Thermo Fisher cat. no. Q332331) and LabChip (Perkin Elmer cat. no. CLS1444006). Samples were then pooled and sequenced on the Illumina NovaSeq 10B paired-end 150 bp flow cell (cat no. 20085594), according to the manufacturer’s recommended protocols. Illumina’s Bcl2fastq2 Conversion software was used to generate de-multiplexed fastq files.

Quality checks were performed on RNA-sequencing data to validate suitability for downstream analysis. Fastq files were checked for quality with FastQC v. 0.12.1^55^, with datasets from all samples having a per-base Phred score greater than 35 across all positions. Reads were aligned to the PAO1 reference genome (RefSeq assembly GCF_000006765.1) with both a splice-aware aligner (STAR v. 2.7.11b^56^) and a non-splice-aware aligner (BWA v. 0.7.18^57^), and quality reports were generated with Qualimap v. 2.3^58^. All reports were collected together for inspection with MultiQC 1.24.1^59^. No prefiltering was performed, as the datasets were high-quality, and downstream quasi-mapping quantification accounts for most quality filtering needs.

Read counts were quantified with Salmon v. 1.10.3.^60^ To do this, the reference genome assembly, CDS, and GFF annotations for *Pseudomonas aeruginosa* str. PAO1 (RefSeq assembly GCF_000006765.1) were collected from NCBI. A decoy reference index was generated with Salmon using the CDS as true reference and the *P. aeruginosa* genome as background to screen out potential artifacts. Fastq reads were quantified against this reference index with Salmon in a quasi-mapping fashion invoked with sequence-specific, GC composition, and positional (5’ or 3’) bias correction. Transcript ID to gene ID, gene names, and other gene metadata were extracted from the reference files obtained from NCBI and prepared as a mapping table. Computation was performed with the help of GNU Parallel v. 20210822^61^ over the UM Advanced Research Computing high performance computing clusters.

The R package DESeq2 v. 1.44.0 was used for differential gene expression analysis, clusterProfiler v. 4.12.0 was used for overrepresentation and gene set enrichment analyses, and Metaboanalyst pathway module was used to identify metabolic pathways predicted by metabolite concentrations in injured relative to healthy BALF.

### Prospective human cohort analysis

We performed a prospective cohort analysis of patients (n = 113) at Clínica Universidad de La Sabana in Chía, Colombia, who received mechanical ventilation for at least 48 hours.

Inclusion criteria were 1) age ≥ 18 years, 2) admission to Clínica Universidad de La Sabana’s critical care unit between January 2020 and July 2022, and 3) invasive mechanical ventilation ≥ 48 hours due to non-infectious respiratory failure. Exclusion criteria included diagnosis of acute pulmonary infection (including COVID) on admission, chronic lung disease, referred patients already on mechanical ventilation, mechanical ventilation over 24 hours before enrollment, prior hospitalization, and prior antibiotics within 48 hours.

Demographic data, comorbidities, symptoms, physiological variables, systemic complications, and laboratory reports at the time of BALF collection were recorded and used for downstream analysis. Subsequent VAP diagnosis was based on current clinical guidelines from the Infectious Diseases Society of America and the American Thoracic Society.^62^ Diagnostic criteria included 1) patients on mechanical ventilation for at least 72 hours, 2) a new or progressive radiographic infiltrate, and 3) at least two of the following symptoms: fever (body temperature > 38°C), purulent tracheal secretions, leukocytosis (leukocyte count > 10,000 cells/μL), or leukopenia (leukocyte count < 4,000 cells/μL). VAP diagnosis was confirmed only if at least one known pneumonia-causing respiratory pathogen was isolated from either endotracheal aspirate (> 10^6^ CFU) or BALF (>10^4^ CFU) after more than 48 hours of intubation.

Following patient enrollment and within 12 hours of intubation, BALF was collected via standard mini-BAL procedure. Cellular BALF aliquots were frozen promptly within one hour of collection and stored at −80°C for downstream analysis. Upon sample transfer to the University of Michigan, BALF was thawed and cellular fraction was separated by centrifuging at 16,000 x *g* for 30 minutes. Supernatant was collected and stored at −80°C for subsequent analysis via IgM ELISA (described under *Lung injury and inflammation*) and spectrophotometric *ex vivo* bacterial growth studies (described under *In vitro and ex vivo bacterial culture*).

### Human BALF untargeted metabolomics

We performed a metabolomics analysis of human BALF on a subset of patients with sufficient sample volume (n = 71) to measure metabolites in the airspace of intubated patients. Sample preparation for the assay has been previously reported.^40^ Briefly, stored BALF samples were thawed on ice and centrifuged (16,000 rpm for 10 min at 4°C). The supernatant was collected and cold (−20°C) methanol:chloroform (9:1; 400 µL) was added to each 100 µL of BALF supernatant and shaken for 5 min at room temperature. Samples were then centrifuged (14,000 rpm for 15 min at 4°C) and supernatants (20µL) were dried (90 min, 35°C) after which O-methoxyamine in pyridine (10 µL of 15 mg/mL) was added. Samples were then vortexed for 10 min and incubated in the dark at room temperature for 16 hours. Subsequently, *N,O-* bistrifluoroacetamide with 1% trimethylchlorosilane (10 µL) was added and the samples were incubated (70°C for 1 h). They were then cooled to room temperature. Methyl stearate in heptane (10 mg/L; 60 µL) was added as an internal standard and the samples were vortexed (10 min).^63–65^

The metabolomics assay was performed using an Agilent Technologies Gas Chromatography 7890B GC system coupled to a Time-of-Flight Mass Spectrometer QTOF 7250 (Agilent Technologies, Waldbronn, Germany). The derivatized sample (1 µL) was injected onto a HP-5MS column (30 m x 0.25 mm x 0.25 µm), using helium as the carrier gas at a constant gas flow (0.7 ml /min). The injector temperature was set to 250°C and split ratio was 30:1. The gradient temperature program started at 60°C for 1 min, then increased to 325°C at a rate of 10°C/min. The GC-MS transfer line was set at 280°C, filament source at 250°C and quadrupole at 150°C. The electron ionization source was set at 70 eV and the mass spectrometer was operated in full scan mode from 50 to 600 m/z at a scan rate of 5 scans/s. Quality control (QC) samples were prepared by pooling equal volumes from each BALF sample. A randomly selected QC sample was injected into the system every 10 samples to assess instrument drift. During data analysis, MVA based on principal component analysis (PCA) was applied to evaluate the quality of the acquired data, and to verify that the QC samples clustered together in these models to demonstrate the stability of the analytical system.

Spectral identification of metabolites started with deconvolution of total ion chromatograms using Agilent Unknowns Analysis B.10.0 and alignment was done using Agilent MassProfiler Professional. The retention time and the mass spectrum of features were compared with the Fiehn GC-MS Metabolomics Retention Time Locked (RTL) Library.^66^ It is essential to note that the level of identification achieved for this platform using Metabolomics Standards Initiative (MSI) criteria was 1, which includes MS spectra and retention time (RT) confirmation. Based on the comparison, annotation levels were reported according to the MSI.^67^ The processed data was exported to the Agilent MassHunter Quantitative software for integration and the areas of each metabolite were normalized according to the response of the QCs using the free software SERRF (https://slfan.shinyapps.io/ShinySERRF/). Only named metabolite features present in at least 80% of the samples and with a coefficient of variation less than 30% in the QC samples were considered for statistical analysis.

### Data analysis

Hypothesis tests used for analyses are indicated in the figure legends for all panels containing significance stars. All statistical tests used *p* = 0.05 as the threshold for significance, except for transcriptomics analysis, where a more conservative threshold of *p* < 0.01 was used. Significance ranges are indicated in all figures as follows: ns = *p* ≥ 0.05; * = *p* ≤ 0.05; ** = *p* ≤ 0.01; *** = *p* ≤ 0.001; **** = *p* ≤ 0.0001.

## Data Availability

Transcriptomic sequencing data have been deposited to NCBI’S Gene Expression Omnibus database^68^ (GEO series accession: GSE278542).

## Acknowledgements

This research was supported in part by the University of Michigan Germ-Free Core, the University of Michigan Biochemical NMR Core, the University of Michigan Advanced Genomics Core, Unisabana Center for Translational Science, and the Universidad de Los Andes Metabolomics Core Facility. The authors thank Larisa Yeomans for her expertise regarding analysis of BALF via NMR spectroscopy and Rishi Chanderraj for helpful discussion of topics related to this manuscript.

## Extended Data Tables and Figures

**Extended Data Table 1:** Retrospective human cohort characteristics. Values indicate count with percentages indicated in parentheses unless otherwise stated. IQR = interquartile range; SD = standard deviation.

| Characteristic | No ARDS<br>(n = 525) | ARDS<br>(n = 258) | Total<br>(n = 783) |
| --- | --- | --- | --- |
| Age, median (IQR) | 60 (49 - 70) | 59 (45 - 68) | 60 (48 - 69) |
| Male sex | 329 (62.7) | 157 (60.9) | 486 (62.1) |
| Race |  |  |  |
| Caucasian | 416 (79.2) | 213 (82.6) | 629 (80.3) |
| African-American | 69 (13.1) | 29 (11.4) | 98 (12.5) |
| Asian | 10 (1.9) | 4 (1.6) | 14 (1.8) |
| Native Indian or Pacific Islander | 0 (0.0) | 3 (1.2) | 3 (0.4) |
| Other | 20 (3.8) | 3 (1.2) | 23 (2.9) |
| Unknown | 10 (1.9) | 6 (2.3) | 16 (2.0) |
| Ethnicity |  |  |  |
| Hispanic or Latino | 18 (3.4) | 4 (1.6) | 22 (2.8) |
| Non-Hispanic or Latino | 495 (94.3) | 246 (95.4) | 741 (94.6) |
| Unknown | 12 (2.3) | 8 (3.1) | 20 (2.6) |
| Hospital unit |  |  |  |
| Medical ICU | 264 (50.3) | 194 (75.2) | 458 (58.5) |
| Surgical ICU | 99 (18.9) | 30 (11.6) | 129 (16.5) |
| Cardiac ICU | 79 (15.1) | 10 (3.9) | 89 (11.4) |
| Burn ICU | 76 (14.5) | 22 (8.5) | 98 (12.5) |
| Moderate Care | 7 (1.3) | 2 (0.8) | 9 (1.1) |
| Maximum SOFA score, mean (SD) | 6.7 (2.3) | 7.6 (2.7) | 7 (2.5) |
| Charlson comorbidity index, median (IQR) | 2 (1 - 5) | 3 (1 - 5) | 2 (1 - 5) |
| Ventilator-free days, median (IQR) | 23 (21-25) | 22 (19-24) | 23 (20 - 25) |
| Length of stay, median d (IQR) | 15.35 (9.0 - 25.6) | 15.84 (10.1 - 26.6) | 15.71 (9.2 - 26.0) |
| 30-day mortality | 134 (25.5) | 89 (34.5) | 223 (28.5) |

**Extended Data Table 2:** Identity of organisms cultured from lower respiratory tract samples from patients in the retrospective observational cohort. Values indicate count with percentages indicated in parentheses unless otherwise stated.

| **Pathogen** | **No ARDS**<br>**(n = 525)** | **ARDS**<br>**(n = 258)** | **Total**<br>**(n = 783)** |
| --- | --- | --- | --- |
| *Pseudomonas aeruginosa* | 24 (4.6) | 19 (7.4) | 43 (5.5) |
| *Staphylococcus aureus* | 25 (4.8) | 11 (4.3) | 36 (4.6) |
| *Stenotrophomonas maltophilia* complex | 5 (1.0) | 2 (0.8) | 7 (0.9) |
| *Escherichia coli* | 5 (1.0) | 3 (1.2) | 8 (1.0) |
| *Enterobacter cloacae* complex | 4 (0.8) | 2 (0.8) | 6 (0.8) |
| *Haemophilus influenzae* | 3 (0.6) | 1 (0.4) | 4 (0.5) |
| *Klebsiella pneumoniae* | 2 (0.4) | 3 (1.2) | 5 (0.6) |
| *Acinetobacter baumannii* complex | 2 (0.4) | 0 (0.0) | 2 (0.3) |
| *Enterobacter aerogenes* | 2 (0.4) | 0 (0.0) | 2 (0.3) |
| Beta-hemolytic, non-Group A *Streptococcus* | 1 (0.2) | 0 (0.0) | 1 (0.1) |
| *Serratia marcescens* | 1 (0.2) | 2 (0.8) | 4 (0.5) |
| *Aspergillus fumigatus* | 1 (0.2) | 1 (0.4) | 2 (0.3) |
| *Pseudomonas* spp. | 1 (0.2) | 0 (0.0) | 1 (0.1) |
| *Pseudomonas fluorescens* group | 1 (0.2) | 1 (0.4) | 2 (0.3) |
| *Providencia rettgeri* | 1 (0.2) | 0 (0.0) | 1 (0.1) |
| *Klebsiella varicola* | 1 (0.2) | 0 (0.0) | 1 (0.1) |
| *Achromobacter* spp. | 1 (0.2) | 1 (0.4) | 2 (0.3) |
| *Rhizopus* spp. | 0 (0.0) | 1 (0.4) | 1 (0.1) |
| *Acinetobacter* spp. | 0 (0.0) | 1 (0.4) | 1 (0.1) |
| Unknown | 1 (0.2) | 0 (0.0) | 1 (0.1) |
| **Total positive cultures** | **81 (15.4)** | **48 (18.6)** | **129 (16.5)** |

**Extended Data Table 3:** Prospective human cohort characteristics. Results are percentages unless otherwise stated. IQR = interquartile range. Anti-pseudomonal coverage indicates receipt of one or more doses of piperacillin/tazobactam, gentamicin, cefepime, or meropenem within 24 hours of intubation.

| **Characteristic** | **P/F > 300**<br>**(n = 34)** | **P/F ≤ 300**<br>**(n = 79)** | **Total**<br>**(n=113)** |
| --- | --- | --- | --- |
| Age, mean (SD) | 49.2 (23.6) | 48.9 (19.9) | 49.0 (21.0) |
| Male sex | 22 (64.7%) | 53 (67.1%) | 75 (66.4%) |
| SOFA score, mean (SD) | 7.3 (2.5) | 8.1 (2.1) | 7.8 (2.2) |
| VAP diagnosis within 7 days of intubation | 6 (17.6%) | 29 (36.7%) | 35 (31.0%) |
| Any antibiotics within 24 h of intubation | 11 (32.4%) | 33 (41.8%) | 44 (38.9%) |
| Anti-pseudomonal antibiotics within 24h of intubation | 4 (11.8%) | 11 (13.9%) | 15 (13.3%) |
| PaO<sub>2</sub>/FiO<sub>2</sub> ratio, mmHg |  |  |  |
| > 300 | 34 | 0 | 34 (30.1%) |
| 200 ≥ 300 | 0 | 35 | 35 (31.0%) |
| 100 ≥ 200 | 0 | 14 | 14 (12.3%) |
| < 100 | 0 | 4 | 4 (3.5%) |
| Alveolar IgM (ng/mL) |  |  |  |
| Duration of mechanical ventilation, median d (IQR) | 3 (3) | 5 (6) | 4 (5) |
| Hospital length of stay, median d (IQR) | 14 (14.5) | 13 (24.5) | 13 (23) |
| In-hospital mortality | 7 (20.6%) | 26 (32.9%) | 33 (29.2%) |

## Extended Data Figures

**Extended Data Fig. 1:**
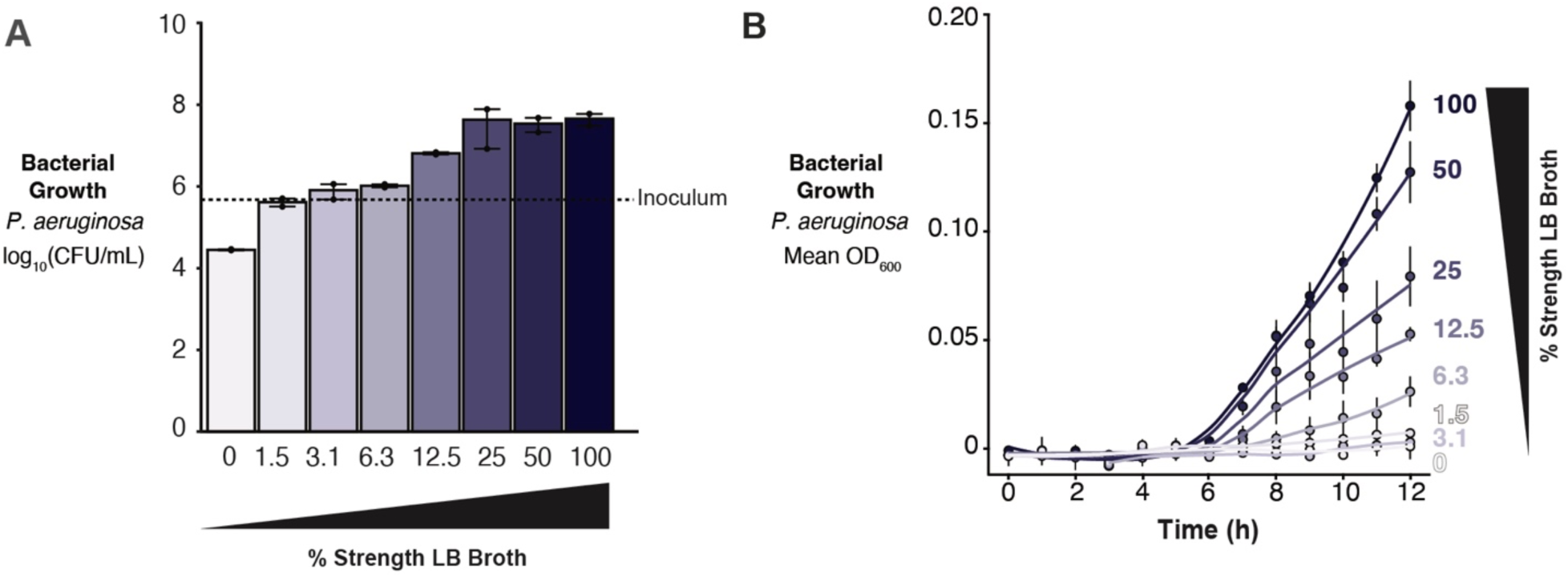
Standard bacterial growth assays can detect differences in nutrient availability. **a,b,** Serial dilution of LB broth, a rich bacterial medium, demonstrates the capacity of (**a**) quantitative CFU plating and (**b**) spectrophotometry to capture CFU changes in *P. aeruginosa* attributable to changes in nutrient availability.

**Extended Data Fig. 2:**
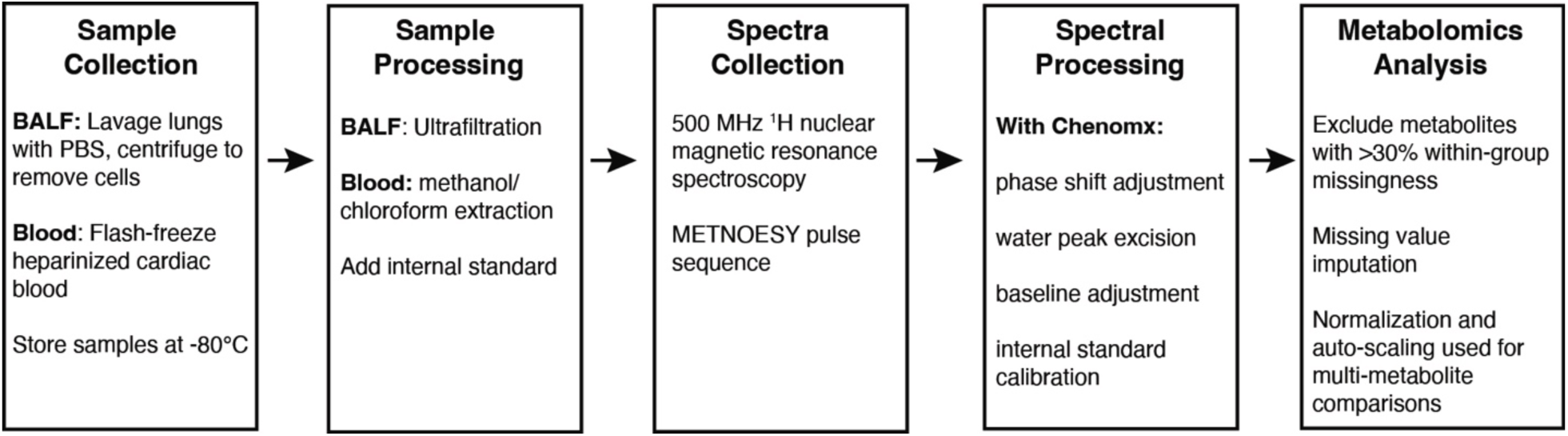
NMR metabolomics workflow used to characterize the murine BALF and whole blood metabolite profiles. Descriptions preceded by BALF or blood indicate a step in the workflow that applies only to the sample type indicated; all other steps are common to both sample types.

**Extended Data Fig. 3:**
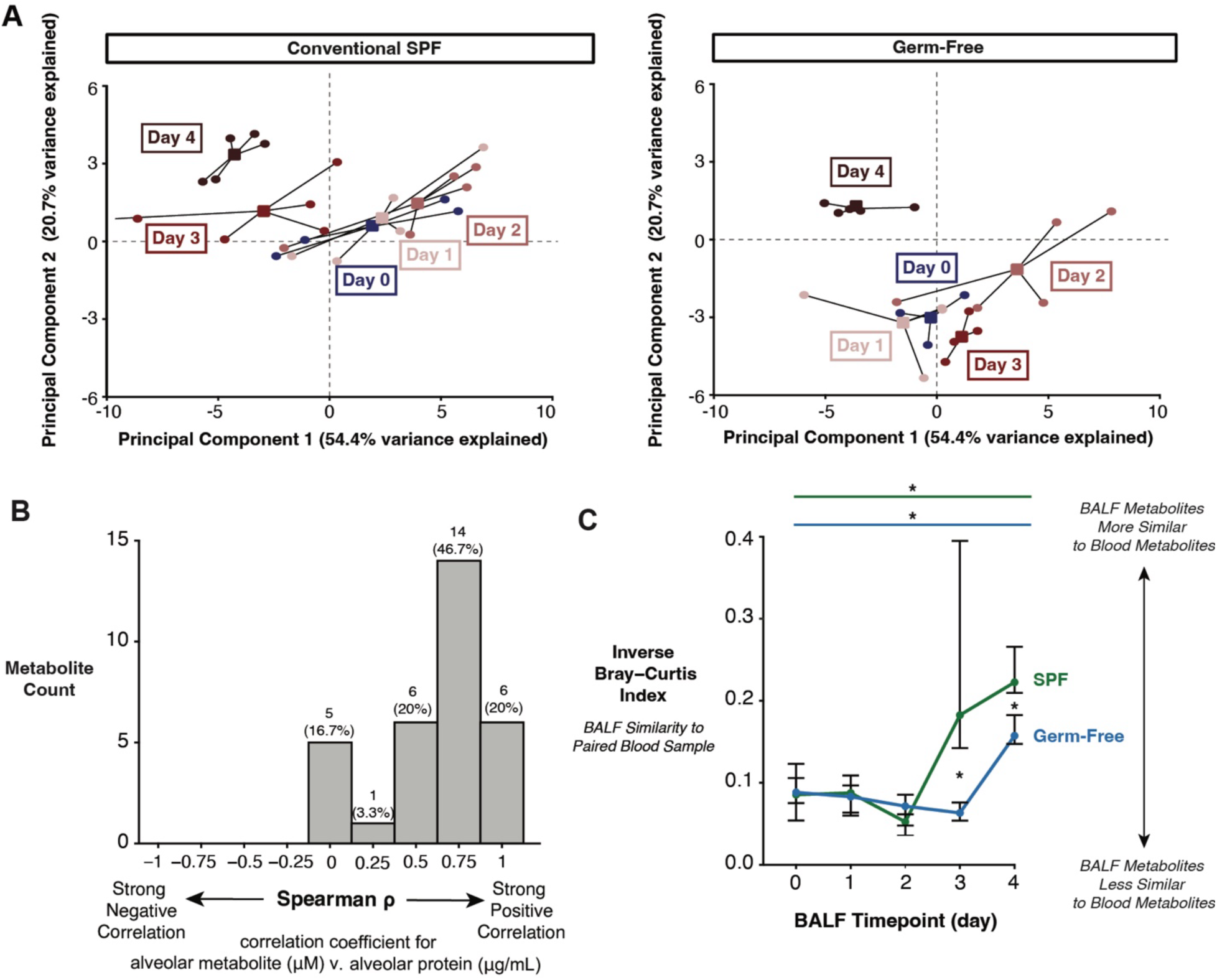
Confirmatory evidence of pulmonary edema as a source of metabolite enrichment in BALF from mice with ALI. **a,** Principal component analysis of the BALF metabolite profiles of conventional and germ-free mice over a 4-day timecourse of hyperoxia exposure. Each point represents an individual mouse and squares represent the centroid of the BALF metabolite profiles; both are grouped and colored by day. **b**, Histogram of Spearman ρ coefficients for BALF total protein concentration with concentrations of each BALF metabolite detected via NMR. **c,** Inverse Bray-Curtis similarity index for metabolites detected in paired BALF and blood samples from conventional and germ-free mice over a 4-day timecourse of hyperoxia exposure. Dots represent median and bars represent IQR. Across-group significance bars represent the results of Kruskal-Wallis rank sum tests, colored by microbiome group. Between-group significance stars represent the results of between-group Wilcoxon rank sum tests. Significance key: ns *p* > 0.05; * *p* ≤ 0.05; ** *p* ≤ 0.01; *** *p* ≤ 0.001; **** *p* ≤ 0.0001.

**Extended Data Fig. 4:**
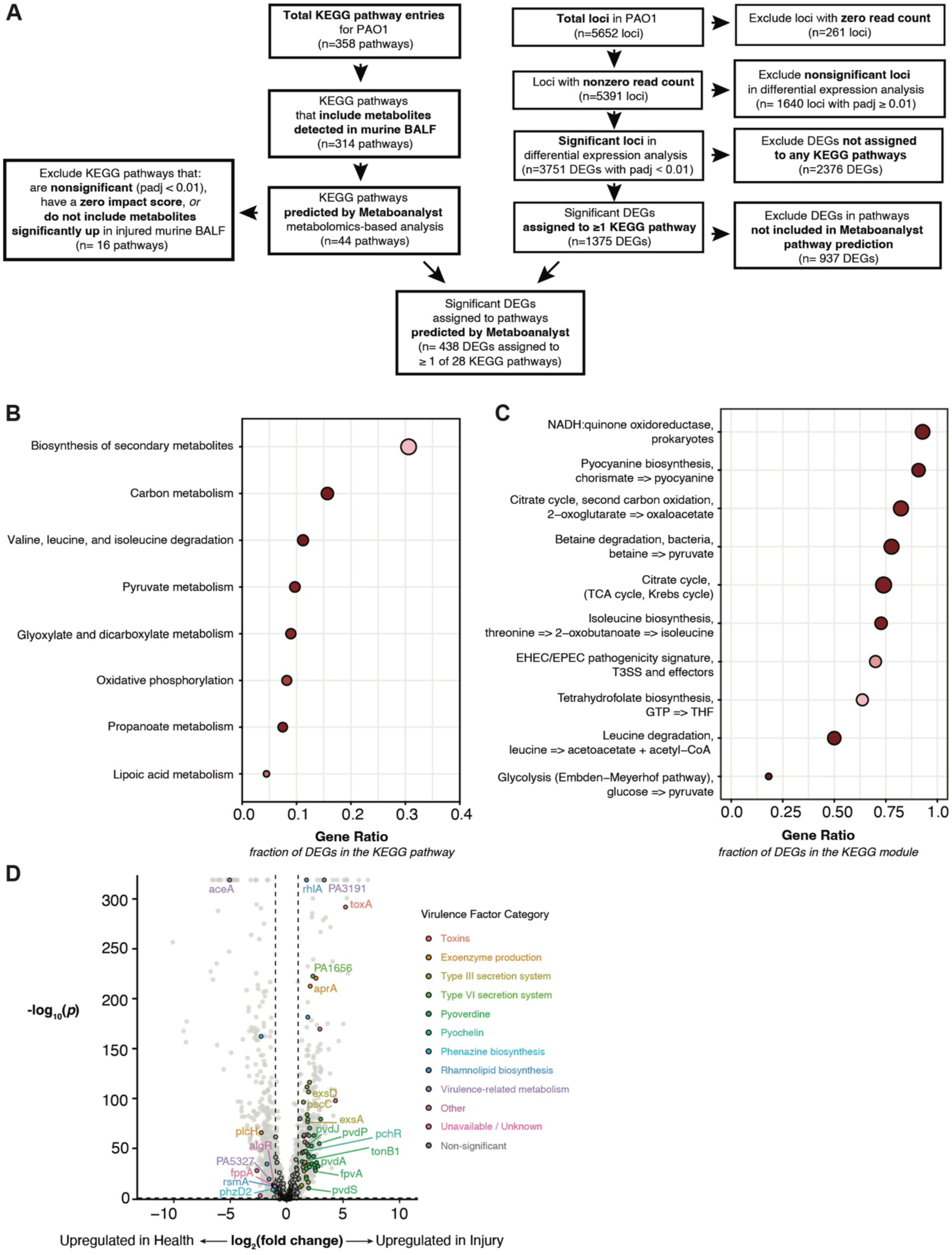
*P. aeruginosa* metabolism and virulence pathways of interest during culture in BALF from mice with and without ALI. **a,** Flow diagram of transcriptomics metabolic pathway analysis. **b,c,** Significant results from overrepresentation analysis using KEGG pathways (**b**) and gene set enrichment analysis using KEGG modules (**c**) of the *P. aeruginosa* transcriptome when cultured in BALF from mice with ALI relative to culture in BALF from healthy mice. Dot size represents gene count and dot color represents adjusted *p*-value. **d,** Volcano plot of virulence genes differentially expressed by *P. aeruginosa* when cultured in BALF from mice with ALI relative to culture in BALF from healthy mice. Dotted lines represent thresholds for fold-change (≥ 2-fold change in expression, vertical) and significance (FDR-corrected *p* ≤ 0.1, horizontal). Genes that meet or exceed significance and fold-change thresholds are colored and selected genes of interest are labeled. Significance key: ns *p* > 0.05; * *p* ≤ 0.05; ** *p* ≤ 0.01; *** *p* ≤ 0.001; **** *p* ≤ 0.0001.

**Extended Data Figure 5:**
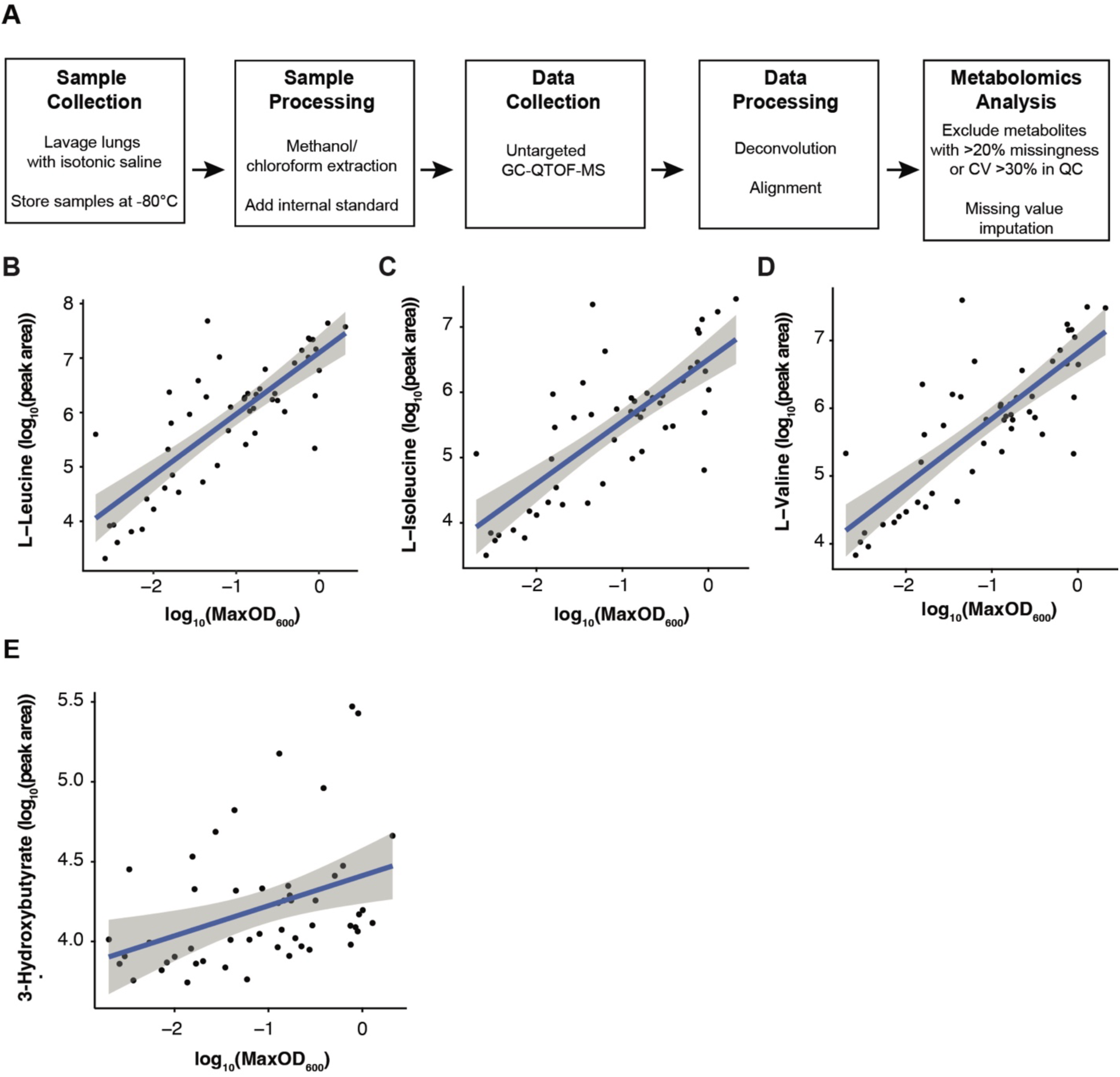
Metabolomic profiling of BALF samples from intubated patients. **a,** GC-QTOF-MS metabolomics workflow used to characterize human BALF metabolite profiles. **b-e**, Correlation of metabolites detected in human BALF via GC-QTOF-MS with paired maximum optical density of *P. aeruginosa* cultured in paired BALF aliquots, including leucine (**b**), isoleucine (**c**), valine (**d**), and 3-hydroxybutyrate (**e**). Blue lines with gray shading indicate linear model with 95% confidence interval.

